# STRUMP-I: Structure-based machine learning approach to pMHC-I binding prediction using force field energy features

**DOI:** 10.1101/2025.09.03.674126

**Authors:** Adam Voshall, Jeongjun Chae, Honglan Li, Junsu Ko, Woongyang Park, Eunjung Alice Lee, Yoonjoo Choi

## Abstract

The adaptive immune system monitors cellular integrity by recognizing short peptides from intracellular proteins presented on Major Histocompatibility Complex class I (MHC-I) molecules, collectively termed peptide-MHC complexes (pMHC), enabling detection of foreign or mutated proteins. With the rising importance of immunotherapies targeting neoantigens in cancers, the ability to accurately predict which peptides will bind to the diverse population of MHC alleles is critically important. Current computational methods for pMHC-I prediction fall broadly into sequence-based methods, which rely heavily on large training datasets, and structure-based methods that leverage structural modeling and energetics of pMHC binding. While sequence-based methods have been popularly used, their performance is dependent on the size and quality of training data. On the other hands, while structure-based approaches can generalize better across diverse MHC alleles, they traditionally depend on identifying a single global minimum energy conformation, an assumption that often fails due to the inherent binding promiscuity of MHC-I molecules. To address these limitations, we developed a STRUMP-I (STRUcture-based pMHC Prediction (for class I)), a novel pMHC binding prediction tool that directly leverages a broad set of force-field-derived energy terms as machine-learning features. STRUMP-I achieves performance comparable to state-of-the-art sequence-based models while significantly outperforming them on MHC alleles with limited representation in training data. Furthermore, STRUMP-I demonstrates strong synergy when integrated with sequence-based methods, notably enhancing prediction precision. The robustness and generalizability of STRUMP-I were confirmed by evaluating its predictive performance on independent, previously unseen datasets, including an experimentally validated cancer neoantigen dataset. This combined approach advances our capability to reliably identify clinically relevant neoantigen targets. The source code and trained models are available at https://github.com/yoonjoolab/STRUMP-I

## Introduction

The Major Histocompatibility Complex (MHC) forms a complex with short peptides derived from endogenously expressed proteins, together known as the pMHC-I, which allows the adaptive immune system to monitor the internal functioning of a cell and surveil for foreign or self-proteins (1). The process involves proteasomal degradation, peptide transport, and loading into the MHC-I, which are then inspected by CD8+ T cells, leading to the proliferation of T cells that can bind to the peptide and the death of the cell presenting it. With the rising importance of immunotherapies targeting neoantigens in cancers (2) and the growing awareness of the importance of T cell immunity in vaccine design (3), accurately predicting which peptides will invoke an immune response is critical. Predicting the MHC-I binding of the peptide forms the core part of many neoantigen prediction pipelines (4). However, as MHC-I is the most hypermorphic gene (5) and the number of known alleles continuously grow (6), this task remains extremely challenging across the vast diversity of possible MHC-I alleles. To address this, significant computational efforts have been made to develop and refine prediction models that can effectively navigate the immense diversity of pMHC-I complexes.

Computational methods for predicting pMHC interactions fall into two broad categories: sequence-based and structure-based methods. In general, sequence-based methods have been widely used in practice since they are faster and more accurate than structure-based ones (7). Nealy all the sequence-based methods recently developed use machine learning techniques to score the strength of the binding directly from the peptide and MHC sequences (8). These methods often incorporate (pseudo-)sequence embeddings that represent key regions of the MHC molecules interacting with peptides. By leveraging large data sets of binding affinity and eluted ligand for the most common MHC alleles, the methods infer residue constraints at specific positions in the peptides and identify biologically meaningful motifs that interact with the binding pockets within the MHC binding groove (7). However, the vast majority of MHC-I alleles have few or no known peptides to train on, limiting their generalizability for vaccine and cancer immunotherapy development (9). To overcome this limitation, the current state of the art methods, including NetMHCpan (10, 11), MHCflurry (12), and HLAthena (13), specifically trained pan-allele models capable of predicting binding scores for MHC alleles absent from the training set, but the performance of these models on low abundance and unseen alleles still lags behind those alleles with better representation (14).

On the other hand, structure-based methods have been underdeveloped compared to sequence-based ones. These methods have primarily focused on accurately reconstructing the orientation and position of the bound peptide within the pMHC binding groove, referred to as the peptide binding mode (15). While these methods typically optimize for binding energy, the scoring functions used in protein design methods such as Rosetta (16) act as correlates for binding affinity. Such approaches also allow the incorporation of modified or non-canonical residues within the peptide (17). While these methods have underperformed the current state of the art sequence-based methods (18), they are less dependent on quality and quantity of the training data and generalize better to MHC alleles and peptides that are poorly represented or missing from the training set (19). Recent methods have reduced the predictive performance gap by either using a per position rather than global scoring scheme (20) or by leveraging the advancements made in *ab initio* protein folding using deep learning by training AlphaFold (21, 22) to learn both structure and binding affinity of the pMHC (19, 23).

Structure-based methods typically rely on a general energy calculation based on the assumption that the pMHC binding happens in the minimum energy state. However, the promiscuity nature of pMHC binding indicates that this assumption is not always met. Considering this suboptimality of the pMHC binding energy landscape, we propose optimizing multiple energy terms of the system to improve prediction accuracy.

Here, we present a STRUcture-based pMHC Prediction (for class I) (STRUMP-I), a novel pMHC binding prediction tool that combines structural modelling with a machine learning-based classifier to discriminate binders from non-binders. Based on the advancement of structural modeling and the idea that the pMHC complex is in general structurally invariable and the pMHC binding classification by atomic interactions can be less data-dependent than sequence-based methods, we directly trained on force-field energy terms for pMHC-I binding interactions. STRUMP-I has comparable performance to the state-of-the-art sequence-based methods but generalizes better to alleles with low or imbalanced representation in the training data. To address computational cost concerns, we also demonstrate how STRUMP-I can be used in combination with the sequence-based methods to improve the Precision of binding predictions.

## Results

### Development of the STRUMP-I model

In order to leverage the structural information about pMHC binding, we developed a machine-learning classifier method, STRUMP-I, that utilizes force-field energy values as features calculated from protein homology-model structure. An overview of the STRUMP-I workflow is presented in Fig. 1a. Briefly, for a given peptide length and MHC type, STRUMP-I identifies the most similar MHC template structure from the known structure database, and a model structure is built by *in silico* mutating differing positions. The mutated structure is then backbone-relaxed and optimized for the AMBER99sb forcefield (26) with Head-Gordon’s Hydrophobic Potential of Mean Force with the Generalized Born model (GB-HPMF) (27) using the Tinker molecular dynamics package (28) to remove potential steric clashes due to the mutations (Fig. 1b). After the relaxation, sidechains were optimized with the FoldX forcefield (29) and their energy terms were directly utilized as features. Apart from the force field energy terms, other quantifiable measures such as the sequence similarities between structure template and input sequence for both MHC and peptide were also included in the feature set.

**Figure 1:**
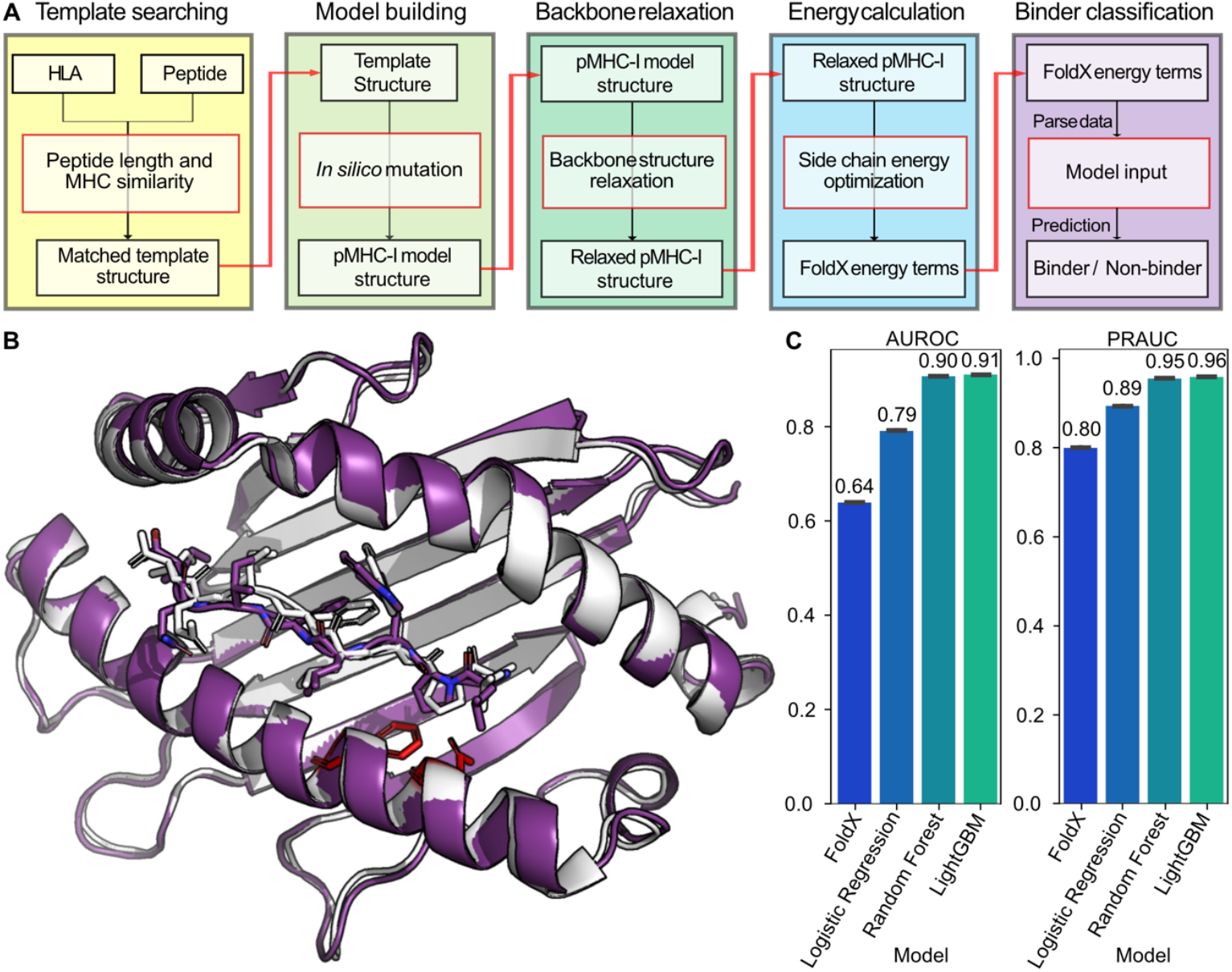
Overview of the STRUMP-I binding prediction pipeline. A) Workflow for the STRUMP-I pipeline from matching the input peptide and MHC-I allele to the experimentally resolved template structures, modifying the selected template to match the inputs, repairing and optimizing the modified structure, and parsing the energetics of the final structure for binding classification. B) Superimposition of the structures for a query sequence with HLA-B*51:01 (white) and matched template sequence with HLA-B*52:01 (purple) after *in silico* mutation and backbone relaxation. Red side chains represent mutations in the MHC. C) AUROC and PRAUC scores for initial tests of different machine learning classifiers on the test set from the IEDB+HLAthena training dataset, showing the average of 100 replicates of 90/10 train/test splits.

To develop the classifier, we built train, validation, and test sets consisting of a combination of the IEDB database and eluted ligand (EL) data used in the development of HLAthena (13). After filtration, this dataset contained 288,269 peptide/MHC-I pairs, consisting of 207,359 binders and 81,110 non-binders (Supplemental File 1). We compared the classification performance using the scaled FoldX interaction energy between the peptide and MHC, which alone is a poor predictor of binding (Figs 1c, S1a) and machine learning algorithms (logistic regression, random forest, gradient boosted machine (GBM), and TabNet neural network (30)) on the full set of features (listed in Table S1) using 100 random seeds with an 90/10 train/test split. We found that the GBM had the highest performance measured by both AUROC and PRAUC on the test data and selected this approach for further development (Fig. 1c). The performance of the GBM was further improved with hyperparameter tuning using Optuna (31). The final model shows clear discrimination between binders (binding affinity < 500nM) and non-binders (binding affinity >= 500nM) (Fig S1b). Based on a SHAP (32, 33) analysis, as expected, many of these important measures were non-bonded energy terms including the overall interaction energy, solvation hydrophobicity and polarity, and electrostatic interactions (Fig. S2). Interestingly, these features also concern the quality of the alignment between MHC alleles or peptides and the templates, in particular the alignment scores and the HLA BLOSUM ratio (the ratio of alignment score to the highest possible score based on a self-to-self alignment to the alignment score for the query and template), indicating that identifying proper structure templates is critical for predicting pMHC-I binding.

### STRUMP-I shows stable prediction even on alleles with low representation

We evaluated the performance of STRUMP-I in comparison to three widely used sequenced-based pMHC binding prediction tools (NetMHCpan v4.1, MHCflurry v2.0, and HLAthena, each with two difference criteria for distinguishing binding) and with the AlphaFold-based peptide binding prediction method (AlphaFold-FineTune (AF-FT)). We first evaluated the performance of these tools using the test set constructed from the IEDB (34) plus HLAthena datasets (13) (IEDB+HLAthena), which consists of 38,302 binders and 9,061 nonbinders from 115 alleles. Within this dataset, STRUMP-I recovered the second highest number of binders (35,149) behind only MHCflurry (36,275 or 35,480, depending on binding measure used) while retaining up to half as many non-binders (2,056 vs. 3,928 for MHCflurry affinity percentile), giving STRUMP-I an overall performance very comparable to the sequence-based methods with a PRAUC of 0.984 and Matthew’s Correlation Coefficient (MCC) of 0.756, compared to a PRAUC of 0.982 and MCC of 0.781 for MHCflurry, a PRAUC of 0.986 and MCC of 0.778 for HLAthena, and a PRAUC of 0.974 and MCC of 0.603 for NetMHCpan (Fig. 2 a-c) with similar scores across other summary metrics (Table S2). AF-FT underperformed on this dataset compared to STRUMP-I and all of the sequence-based methods with a PRAUC of 0.916 and MCC of 0.422. This performance is nearly identical on the validation set used for hyperparameter tuning (Table S3).

**Figure 2:**
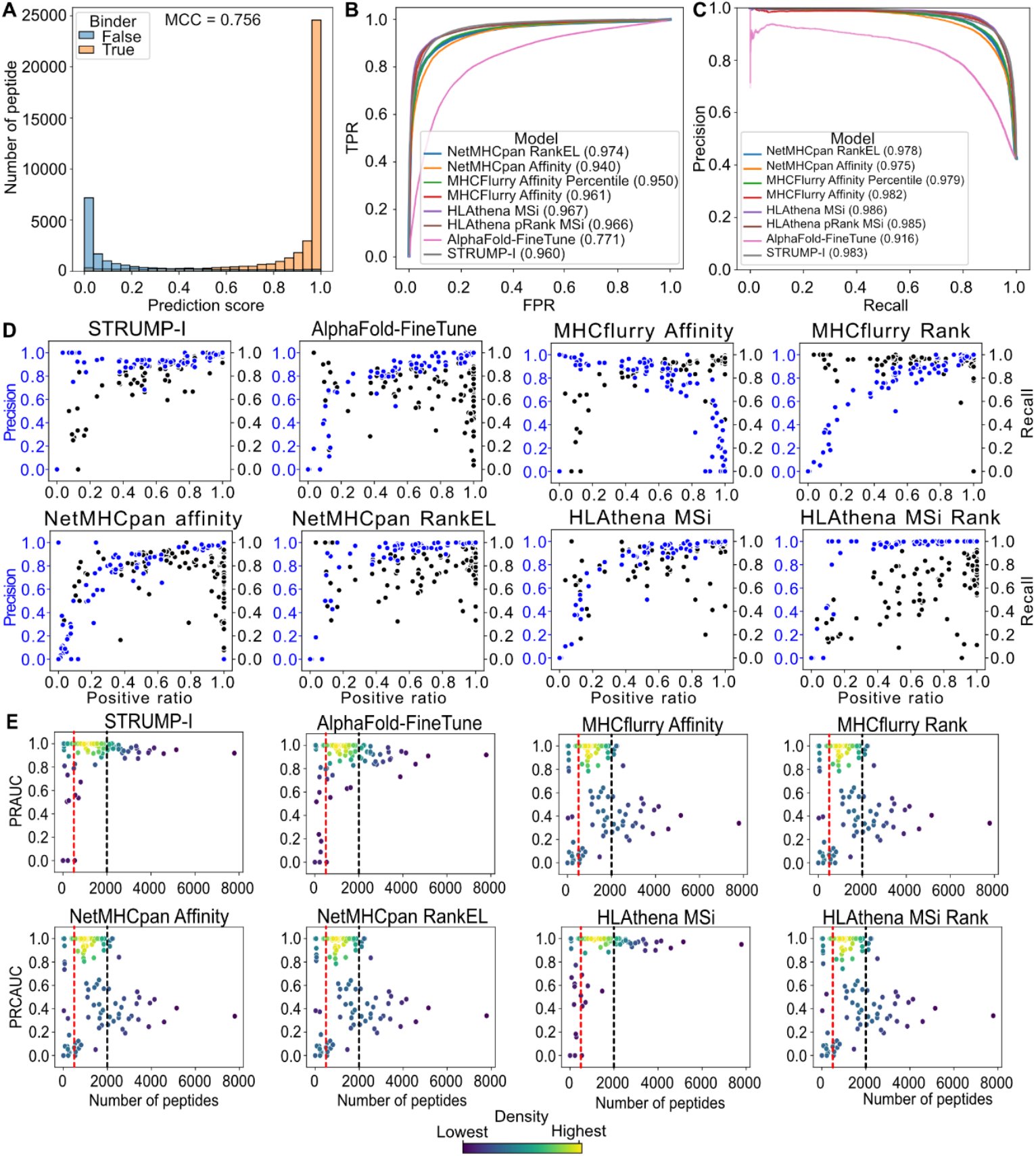
Performance of STRUMP-I on the IEDB+HLAthena test set. A) STRUMP-I prediction scores for binders (orange) and non-binders (blue). B) ROC curves for each of the methods tested on the IEDB+HLAthena test set. TPR: True Positive Rate, FPR: False Positive Rate. C) Precision recall curves for each of the methods tested on the IEDB+HLAthena test set. D) Relationship between positive ratio and precision (blue) or recall (orange). The positive ratio is defined as the ratio between the number of positive and negative peptide/MHC pairs in the training data. E) Scatterplots showing the PRAUC scores for each allele based on the number of peptides present in the IEDB+HLAthena training set, with dots colored by density (blue represent low density, yellow represents high density). Alleles <2,000 peptides in training set (denoted by the black dashed line) constitute the average representation set. Alleles with <500 peptides in the training set (denoted by the red dashed line) constitute the low representation dataset.

These differences in performance are most pronounced for alleles that are less abundant or that have less balanced representation within the training dataset. The number of training peptide/MHC pairs for each allele and positive ratios (number of positive examples divided by the total number of pairs for each allele) are presented in Table S4 and, and the overall distributions for each are shown in Fig. S3. Methods heavily dependent on the underlying data distribution typically show an arch-shaped distribution in performance metrics, characterized by peaks in precision and recall at intermediate positive ratios (balanced representation of positive and negative examples). Performance often drops off at extremes of the positive ratio, highlighting sensitivity to dataset balance. Less data-dependent methods demonstrate relatively stable or level performance across the full range of positive ratios, indicated by the absence of clear peaks or troughs, with precision and recall values distributed evenly from low to high positive ratios. The affinity predictions for NetMHCpan, MHCflurry and MSi score for HLAthena all show this arch-shaped pattern, with a high precision but low recall for alleles with a high positive ratio and a low precision and a high Recall for alleles with a low positive ratio (Fig. 2d). The rank-based scores for each sequence-based method had better precision across positive ratios, but NetMHCpan RankEL and HLAthena MSi Rank in particular had inconsistent recall even for alleles high positive ratios. Only STRUMP-I maintains a high precision across the full range at moderate cost of recall (Fig. 2d).

When the most represented alleles were excluded, and we measured the performance on only alleles with <2000 peptides present in the training set (average representation, Fig. S3), the performance of most tools increased slightly with STRUMP-I increasing to a PRAUC and MCC of 0.995 and 0.828, comparable to HLAthena (PRAUC=0.994, MCC=0.822) and ahead of MHCflurry (PRAUC=0.987, MCC=0.702), NetMHCpan (PRAUC=0.987, MCC=0.581), and AF-FT (F1-score=0.949, MCC=0.359) (Fig. 2e, S2, Table S5). Notably, when we measured the performance on only those alleles with <500 peptides in the training set (low representation, Fig. S3), the performance of all methods, except NetMHCPan using predicted affinity, dropped (Fig. S4, Table S6). This drop in performance was most pronounced in MHCflurry (PRAUC=0.952, MCC=0.732), HLAthena (PRAUC=0.932, MCC=0.5), and AF-FT (PRAUC=0.934, MCC=0.279). In contrast, STRUMP-I’s performance remains consistently high with a PRAUC 0.989 and an MCC of 0.881.

### Benchmarking STRUMP-I with neoantigen datasets

Following benchmarking on the IEDB+HLAthena dataset, we retrained STRUMP-I on the full dataset for benchmarking two additional public neoantigen datasets, TESLA (35) and PRIME (36). These datasets predominantly (PRIME) or entirely (TESLA) consist of average or overrepresented alleles (Table S4). When applied to these additional benchmark datasets, the performance of all of the methods dropped considerably with STRUMP-I’s performance remaining comparable to the sequence-based methods. STRUMP-I’s overall performance on the TESLA dataset with a PRAUC of 0.866 falls between HLAthena and AF-FT on the low end (PRAUCs of 0.759) and NetMHCpan and MHCflurry on the high end (PRAUCs of 0.929 and 0.937) (Fig. 3a, Tables S8 and S9). On the PRIME dataset, STRUMP-I has a lower overall performance than the other methods with a PRAUC of 0.846, with the other methods ranging from 0.953 to 0.988. The differences in performance between STRUMP-I and the other methods is consistent with the differences in performance for the overrepresented alleles (Tables S7, S8, and S9).

**Figure 3:**
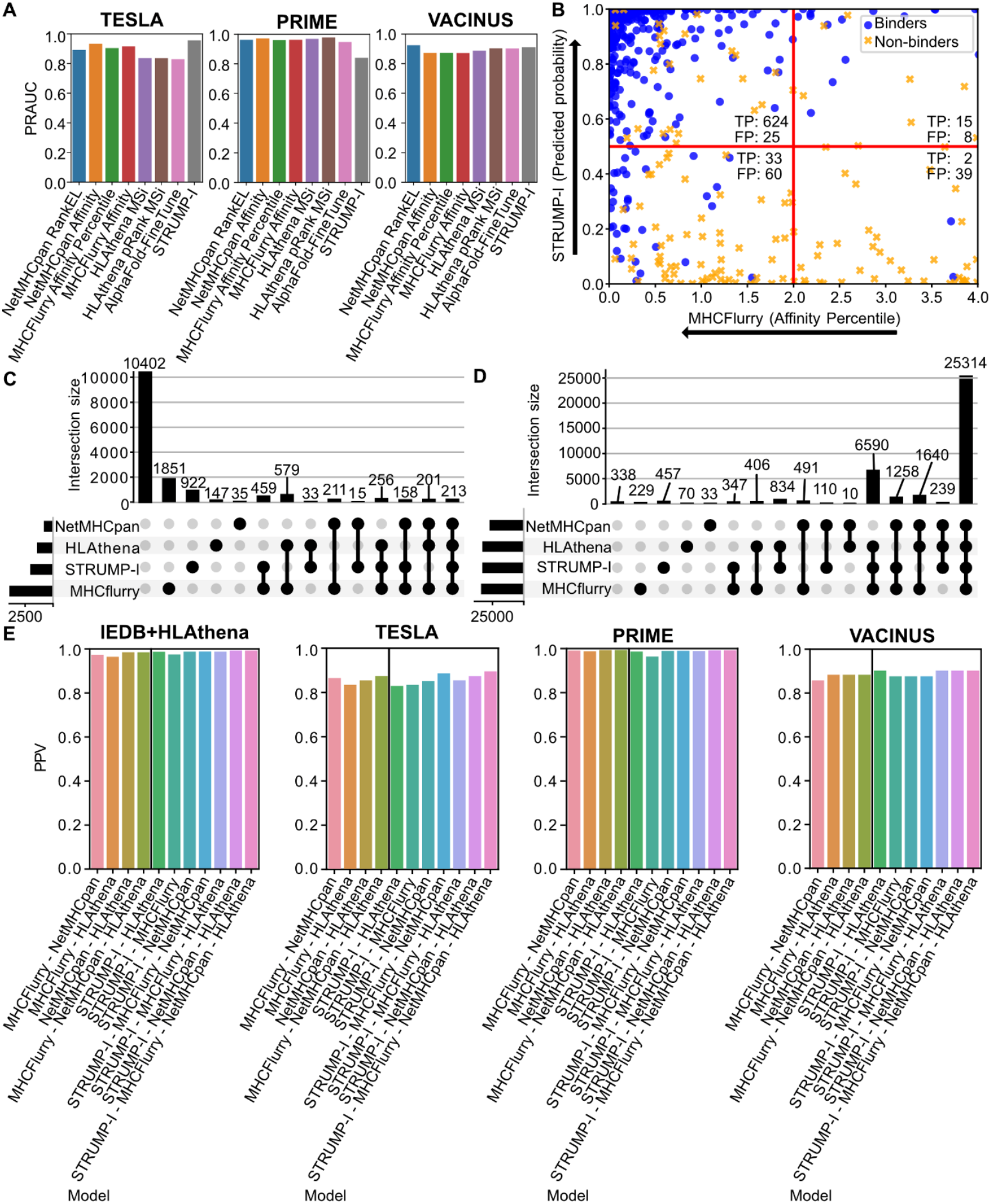
Binding prediction performance on external and VACINUS datasets. A) Bar plots showing the PRAUC scores on the TESLA, PRIME, and VACINUS cancer dataset, respectively. B) Scatter plot showing the Binding Affinity Percentiles from MHCFlurry and the Predicted Binding Probabilities from STRUMP-I for binders (blue dots) and non-binders (orange x’s) for 1,000 peptides from the IEDB+HLAthena dataset. Red lines indicate classification thresholds for each method. Black arrows indicate the direction of higher prediction scores. For MHCFlurry, a lower score indicates the peptide is more likely to bind, while for STRUMP-I a higher score indicates the peptide is more likely to bind. Peptides in the upper left quadrant are predicted to bind by both tools. C) and D) UpSet plots for non-binders (C) or binders (D) predicted as binders by each tool. For quantitative measures and other metrics, see Tables S2, S7, S8, S10. E) Precision scores for the intersection of predictions from multiple tools on each of the benchmark datasets. The left of the vertical bar shows scores for sequence-based methods alone, and the right of the vertical bar shows the scores for sequence-based methods and STRUMP-I.

### Integrating STRUMP-I with sequence-based prediction methods improves precision

In addition to its performance as a standalone binding prediction tool, STRUMP-I can also be integrated with sequenced-based binding prediction methods within neoantigen prediction pipelines to improve the precision of those pipelines. When comparing the prediction scores for true or false positive predictions between the sequence-based methods and STRUMP-I, we found that predominately both tools predict binding for true binders, while false positive predictions are typically only predicted by individual tools (Fig. 3b). While many of the false positive predictions are tool-specific and are filtered out by any combination of tools, the inclusion of STRUMP-I with the sequence-based methods further removes false positive predictions for every combination of sequence-based tools (Figs. 3c and 3d). Consequently, every combination containing STRUMP-I has a higher precision than the corresponding combination without STRUMP-I, reaching as high as 0.991 on the IEDB+HLAthena dataset and 0.990 on the PRIME dataset when combing the predictions of every tool (Fig 3e and Tables S2, S3, S8, S9).

### Prediction of unseen neoantigen candidate peptides

In addition to public datasets, we used the VACINUS neoantigen dataset, a set of 129 neoantigen candidates from 33 cancer patients (6 colorectal cancer, 2 melanoma, 14 hepatocellular carcinoma, and 11 gastric cancer) and spanning 6 overly represented and 1 average represented alleles (Tables 1, S10) (37), using NetChop and MHCflurry to predict protein cleavage sites and pre-filtered the peptides using NetMHCpan and MHCflurry for binding potential before synthesizing the peptides and confirming binding with Immunitrack ApS (Copehagen, Denmark). Based on how this dataset was generated, both NetMHCpan and MHCflurry have a recall of 1, but both also falsely predicted 38 peptides to bind leading to a precision of 0.85 and a PRAUCs ranging from 0.928 to 0.96 (Fig. 3a,e, Table S11). Consistent with the other benchmark datasets, STRUMP-I further reduces the number of false positive predictions in this dataset down to 28 at a moderate cost to recall, increasing the precision to 0.870 (Fig. 3e, Table S11) with a PRAUC of 0.928 (Table S11). This performance makes STRUMP-I the second most precise on this dataset comparable to HLAthena (precision of 0.876 and PRAUC of 0.907), and ahead of AF-FT (precision of 0.864 and PRAUC of 0.905).

## Discussion

Here, we presented STRUMP-I, a novel structure-based peptide/MHC class I binding prediction pipeline, along with the VACINUS dataset consisting of 110 binders and 19 non-binders (37). Previously, sequence-based peptide/MHC class I binding prediction methods have outperformed structure-based prediction methods. With STRUMP-I and AF-FT, structure-based methods have an overall performance very comparable to sequence-based methods. As with the generalizability of peptide binding prediction performance to proteins beyond MHC seen with AF-FT, we find that STRUMP-I generalizes better to underrepresented alleles of MHC-I both in terms of total the number of examples of that allele and for the ratio of positive to negative examples in the training set than the sequence-based methods despite the recent advances in pan-allele training these methods employ. The competitiveness and generalizability of STRUMP-I compared to the sequence-based methods is even more striking given that the validation and test sets built from the IEDB and HLAthena datasets were partial to entirely represented in the training data for the sequence-based methods, giving those methods an advantage over STRUMP-I on these tests. Despite this advantage, only MHCflurry slightly outperformed STRUMP-I overall due to its better performance on the highest represented alleles and fell behind STRUMP-I once those alleles were removed. While this performance dropped slightly when STRUMP-I was applied to the TESLA and PRIME datasets, its performance remained in line with HLAthena. As the PRIME dataset mostly and the TESLA dataset entirely consist of overrepresented alleles with positive ratios around 0.5, this relative drop in performance is consistent with the performance difference on the overrepresented alleles in the test set.

During our analyses, we compared STRUMP-I to another recent structure-based pMHC binding prediction method, AF-FT. STRUMP-I and AF-FT performed comparably on TESLA and PRIME, STRUMP-I significantly outperformed AT-FT on the test data derived from the IEDB and HLAthena datasets. While the computational requirements for STRUMP-I to generate the 3d structure of the pMHC are greater than the sequence-based methods, the improved performance on poorly represented alleles in the training data and generation of the pMHC structures for use in downstream analyses justifies the additional computational time, especially as STRUMP-I can be used as a post-prediction filter for sequence-based tools as part of a neo-antigen prediction pipeline, as applied to the VACINUS cancer dataset. In this use case, the more efficient sequence-based methods can quickly screen a patient’s full proteome for potential binders, then STRUMP-I can filter false positive predictions to create a higher confidence set of candidate peptides before experimental validation.

As a standalone tool, STRUMP-I matches the overall performance of sequence-based tools across multiple datasets. Importantly, by utilizing structural information about the pMHC, it is less dependent on the size and quality of the training data for underrepresented alleles, outperforming the sequence-based methods both on alleles with low representation and a low ratio of positive to negative examples. For well represented alleles, STRUMP-I can instead be integrated with the sequence-based methods to remove false-positive predictions, using the sequence-based methods to quickly screen the full proteome for potential binders then using STRUMP-I to refine the predictions. Moving forward, the structural outputs generated by STRUMP-I present promising opportunities for future research, particularly in integrating structural features such as solvent accessibility, peptide flexibility, and detailed atomic interactions into predictive models for immunogenicity and TCR specificity.

## Materials and Methods

### Datasets used

#### IEDB dataset

All human MHC ligand assay results were exported from IEDB (http://iedb.org) (34) and filtered following the strategy outlined by Sarkizova, *et al*. Only peptide/MHC pairs with a quantitative measurement in dissociation constants were kept. Peptides detected using any of the “purified MHC/direct/radioactivity/dissociation constant KD”, “purified MHC/direct/fluorescence/half maximal effective concentration (EC50)” or “cellular MHC/direct/fluorescence/half maximal effective concentration (EC50)” assay types were removed, along with any peptides that were present multiple times in the database and where the difference between the maximum and minimum value for the log-transformed affinity (1-log(nM)/log(50000)) was > 0.2. For the remaining peptide/MHC-I pairs, peptides with an affinity <= 500nM are considered binders, and those with an affinity > 500nM are considered non-binders. After filtering, there was a total of 115,644 pairs from this dataset.

#### HLAthena datasets

We combined the IEDB dataset with the MHC-I eluted ligands from mono-allelic cell lines published alongside HLAthena (13). Because these peptides were derived from cell lines that expressed only a single MHC allele, all peptides are considered binders for the allele expressed in the cell line they were detected in. We filtered any non-canonical alleles and peptides, leaving a total of 173,142 peptide/MHC-I pairs. After merging the datasets, we removed any redundant peptide/MHC-I pairs. The final IEDB+HLAthena dataset has 288,269, consisting of 207,359 binders and 81,110 non-binders.

#### Public neoantigen datasets

We downloaded the TESLA (35) dataset via the SYNAPSE consortium. As with the IEDB and HLAthena datasets, we considered any peptide with a binding affinity <= 500nM as a binder. The PRIME (36) dataset was taken from the original Schmidt, et al. publication. The peptides from “Random” were considered non-binders, and peptides from all other studies of origin were considered binders. For the immunogenicity predictions, the ground truth was taken from the data sets. The PRIME % rank for the corresponding MHC-I allele was calculated using PRIME2.0 (38).

#### VACINUS neoantigen dataset from cancer patients

The VACINUS neoantigen candidates were collected from 33 cancer patients (6 with colorectal cancer, 2 with melanoma, 14 with hepatocellular carcinoma, and 11 with gastric cancer) recruited at Asan Medical Center (IRB number 2022-0263) (37). All procedures were carried out in accordance with relevant guidelines and regulations. We selected highly expressed nonsynonymous or indel mutations using whole transcriptome sequencing (WTS) data. The selection criteria included VAF > 0, read counts > 10 at the transcript level, and the sum of transcripts per million (TPM) calculated based on genes > log 0.4. We predicted cleavage products using NetChop and MHCflurry. Cleaved peptide products were initially screened using NetMHCpan and MHCflurry. Peptides from the intersection of these prediction sets were experimentally validated using the NeoScreen MHC/Peptide Binding Assays (Immunitrack ApS, Copenhagen, Denmark). The fluorescent peptide–dextramers were synthesized by Immudex, Copenhagen, Denmark. The final dataset consists of 120 peptide/MHC-I pairs, of which 101 are binders and 19 are non-binders.

### STRUMP-I pipeline

#### Generation of model structure and energy calculation

826 template pMHC PDB structures were downloaded from the IMGT database (any (TCR-)pMHC-I complexes). Following the download, we extracted the 181 amino acids comprising the G-domain of the MHC-I, which includes the peptide binding regions. Given query peptide and MHC-I sequences, STRUMP-I first finds pMHC-I complex structures with the same peptide length. Then the most similar MHC-I allele is identified by aligning the query allele to the template sequence using the BLOSUM62 scoring matrix. If there are multiple complexes left, it finds the most similar peptide using the PAM30 scoring matrix. Following template selection, STRUMP-I uses the BuildModel module from FoldX to replace peptide and the non-matching regions of the MHC-I from the template with the query. The mutated PDB is then relaxed with the Tinker molecular dynamics package (version 8.9.5) (28), using the AMBER99sb parameter with the GB-HPMF implicit solvent model. To generate the final model and create energy features, the relaxed model is further optimized using FoldX until the energy is no longer improved. The final energetics of the structure are calculated using the AnalyzeComplex module of FoldX.

#### Model Training and classification

For benchmarking STRUMP-I on the IEDB plus HLAthena dataset, we split the data into train/validate/test sets along an 8:1:1 ratio using Scikit-learn. We built STRUMP-I using the python implementation of version 4.0.0 of LightGBM (39). We tuned the hyperparameters for the model using version 2.3.0 of the Optuna optimization library for python for 10,000 rounds with pruning (31). The hyperparameters used to train the final model are presented in Table S12. Following benchmarking on this dataset, we retraied STRUMP-I using the full dataset and the same hyperparameters to make the final model for benchmarking on the TESLA, PRIME, and the VACINUSs neoantigen dataset. We tested the feature importance for this final model using version 0.43.0 of the SHAP python library (32, 33).

### Model performance comparison

The performance of STRUMP-I was compared against three sequenced-based and one other structure-based binding prediction tools. The three sequence-based tools, NetMHCpan (version 4.1) (10), HLAthena (13), and MHCflurry (version 2.0) (12), were benchmarked using both the binding cutoff score suggested by each article (%Rank EL <= 0.5 for NetMHCpan, prank.MSi <=0.2 for HLAthena, and Affinity percentile <= 2%) along with the predicted affinity scores using the same cutoff of an affinity <= 500nM as for the experimental exalts. For HLAthena, since it does not report a predicted affinity, we instead use an MSi score >= 0.5 as a cutoff. The other structure-based method, AlphaFold-FineTune (AF-FT) (19), was benchmarked in two steps. First, we downloaded all pMHC PDB files from the PDB website (2023.12) and found the most similar template PDBs for each input pMHC pair based on their sequence information. Similarity was calculated using the BLOSUM62 matrix. Next, we ran the AF-FT with default options, resulting in a PAE score per pMHC. We classified pMHCs based on the PAE score using logistic regression, considering a logit score greater than 0 as positive.

## Acknowledgments

This study was supported by the Suh Kyungbae Foundation (E.A.L), DP2 AG072437 (E.A.L), and R01 AG078929 (E.A.L). Y.C. was supported by the Medical Research Center Program (NRF-2020R1A5A2031185), the National Immunotherapy Innovation Center (NRF-2020M3A9G3080281), and the Basic Science Research Program (NRF-2021R1F1A1063769).

## Notes

https://github.com/yoonjoolab/STRUMP-I

### Competing Interest Statement

W.P. is a founder and CEO of Geninus Inc. The other authors declare no competing interests.

