## Supplementary material for "STRUMP-I: Structure-based machine learning approach to pMHC-I binding prediction using force field energy features": S1 File

### Figures

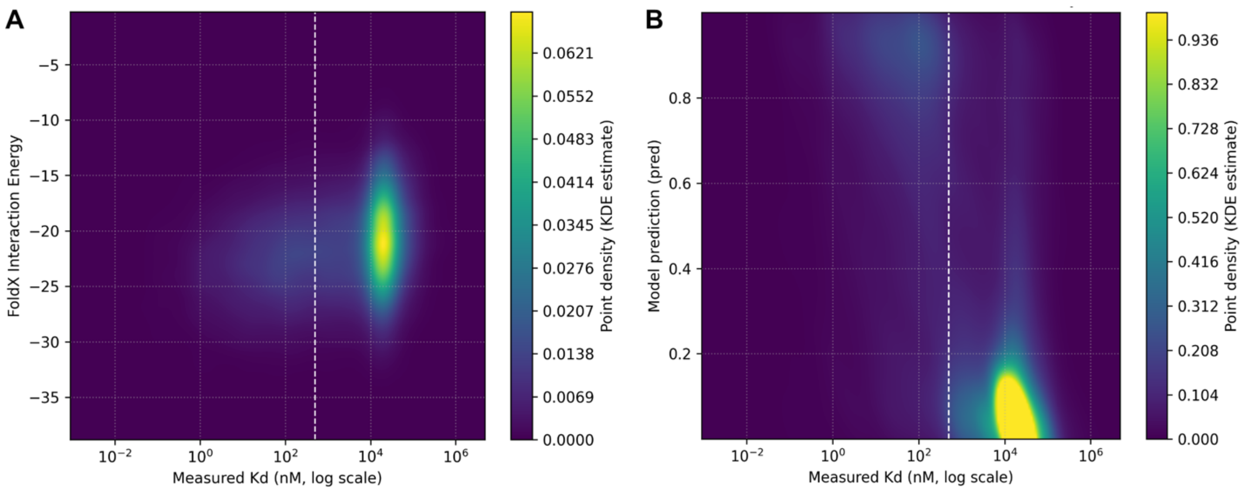


**Figure S1: Density plots of binding affinity in the test set.** White dashed line represents a binding affinity 500 nM used as a threshold to define binders and non-binders. A) Binding affinity and interaction energy calculated by FoldX. B) Binding affinity and prediction score from STRUMP-I.

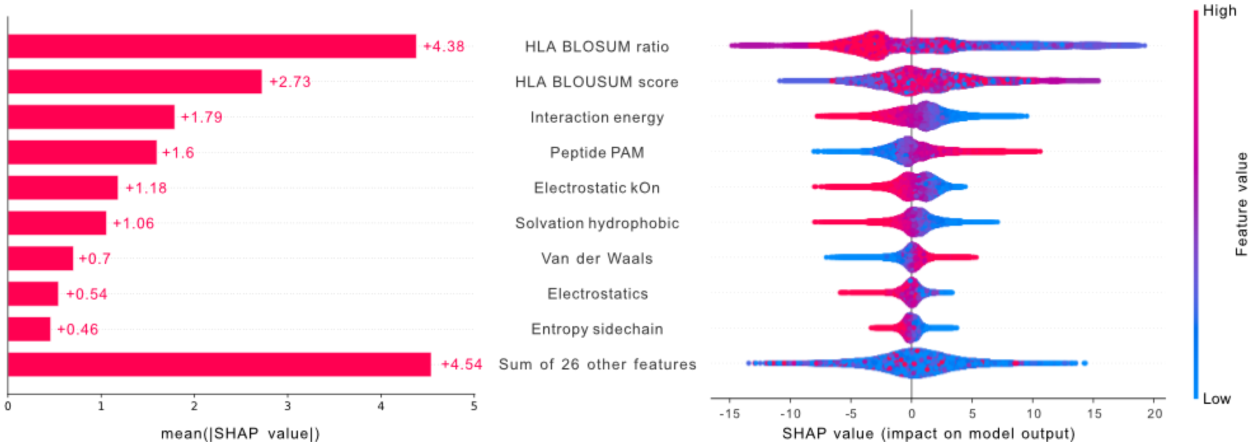


**Figure S2: SHAP summary values for the top used for STRUMP-I binding prediction on the in-house cancer dataset.** For a description of the features, see Table S1. The features are sorted by the magnitude of the SHAP value to highlight the overall most important features driving predictions. Left) Barplot showing the mean absolute value for the relative impact on the model output. Right) Beeswarm plot where red and blue dots indicate peptides with higher or lower than median values for the feature, respectively. The distance from the center line indicates the relative impact of the feature on the final prediction score for that peptide.

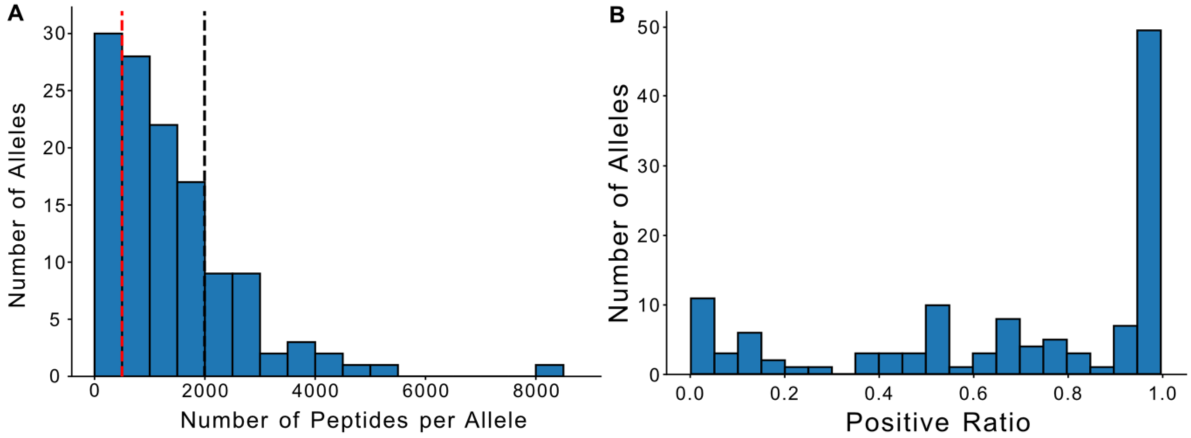


**Figure S3: Histograms for number of examples per allele in the training set.** A) Total number of peptides per allele. Red dashed line – 500 peptides per allele threshold. Alleles to the left of the line are classified as "Low represention". Black dashed line – 2000 peptides per allele. Alleles to the right of the line are classified as "Over represention". Alleles between the lines are classified as "Average representation". B) Positive ratio (percent of positive examples) per allele.

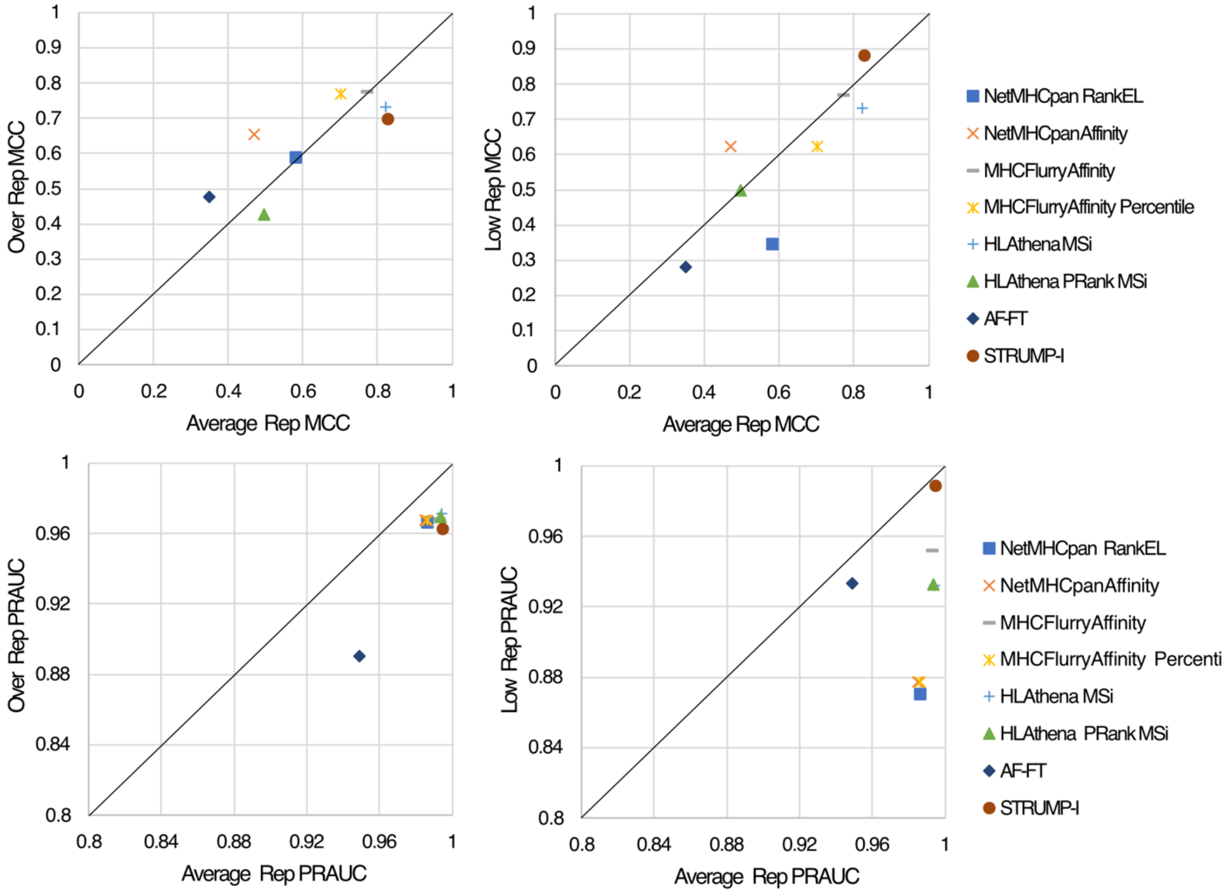


**Figure S4: Method performance on Over, Average, and Low representation alleles.** Performance measured by Matthew's Correlation Coefficient (MCC) (top) and PRAUC (bottom). Over representation are alleles with >= 2,000 pMHC pairs in the training set. Average representation are all alleles with < 2,000 pMHC pairs in the training set. Low representation are all alleles with < 500 pMHC pairs in the training set. The line represents equal performance on alleles in each category. Methods under the line show decreased performance on the over representation or low representation alleles.

**Tables**

**Table S1.** Description of features used to classify binders in STRUMP-I

| Feature | Description |
|---|---|
| HLA BLOSUM score | Alignment score of the input MHC allele to the template using the BLOSUM62 scoring matrix |
| HLA BLOSUM ratio | Ratio of the best alignment score for the input to the maximum possible alignment score for the template sequence |
| Peptide PAM score | Alignment score of the input peptide to the template using the PAM30 scoring matrix |
| Improvement during repair | The amount of energy improvement during iterative PDB repair process |
| Intraclashes Group 1 (MHC) | The Van der Waals' clashes of residues at the interface of the complex with their own molecule |
| Intraclashes Group 2 (peptide) | The Van der Waals' clashes of residues at the interface of the complex with their own molecule |
| Interaction energy | Binding energy |
| Backbone Hbond | The contribution of the backbone hydrogen bonds |
| Sidechain Hbond | The contribution of sidechain-sidechain-backbone hydrogen bonds |
| Van der Waals | The contribution of Van der Waals interactions |
| Electrostatics | Electrostatics interactions |
| Solvation polar | The penalty for burying polar groups |
| Solvation hydrophobic | The contribution of hydrophobic groups |
| Van der Waals clashes | The penalty due to interresidue Van der Waals clashes |
| Entropy sidechain | The entropy cost of fixing the side chain |
| Entropy mainchain | The entropy cost of fixing the main chain |
| Torsional clash | The intraresidue Van der Waals torsional clashes |
| Backbone clash | The backbone-backbone intramolecular Van der Waals clashes |
| Helix dipole | The electrostatic contribution of the helix dipole |
| Electrostatic kon | The electrostatic interaction between molecules in the precomplex |
| Energy Ionisation | The contribution of ionisation energy |
| Stability Group 1 (MHC) | The DG energy term for the MHC protein |
| Stability Group 2 (peptide) | The DG energy term for the peptide |
| Interface Residues | Number of residues in the interface |
| Interface Residues Clashing | Number of residues that are clashing |

| | |
|---|---|
| Interface Residues VdW Clashing | Number of residues with Van der Waals clashing |
| Interface Residues BB Clashing | Number of residues with backbone clashing |
| Water bridge | The contribution of water bridges |
| Disulfide | The contribution of disulfide bonds |
| Cis bond | The cost of having a cis peptide bond |
| Sloop entropy | Loop entropy of the side chain |
| Mloop entropy | Loop entropy of the main chain |
| Final energy | The total energy of the final model |

Table S2. Performance metrics on the test set from training data

| Model | TP | FP | TN | FN | Accuracy | Recall | Specificity | Precision | AUC | PRAUC | F1 | MCC |
|---|---|---|---|---|---|---|---|---|---|---|---|---|
| NetMHCpan RankEL | 29985 | 680 | 14802 | 8381 | 0.8317 | 0.7816 | 0.9561 | 0.9778 | 0.9744 | 0.9784 | 0.8687 | 0.6028 |
| NetMHCpan Affinity | 29095 | 833 | 14649 | 9271 | 0.8124 | 0.7584 | 0.9462 | 0.9722 | 0.94 | 0.9746 | 0.8521 | 0.5668 |
| MHCFlurry Affinity | 35480 | 1810 | 13682 | 2886 | 0.9128 | 0.9248 | 0.8831 | 0.9515 | 0.9612 | 0.9823 | 0.9379 | 0.7813 |
| MHCFlurry Affinity Percentile | 36275 | 3928 | 11554 | 1832 | 0.8925 | 0.9519 | 0.7463 | 0.9023 | 0.95 | 0.9785 | 0.9264 | 0.7509 |
| HLAthena MSi | 35103 | 1429 | 14053 | 3262 | 0.9129 | 0.915 | 0.9077 | 0.9609 | 0.9673 | 0.9861 | 0.9374 | 0.7779 |
| HLAthena prank MSi | 25339 | 176 | 15306 | 13027 | 0.7548 | 0.6605 | 0.9886 | 0.9931 | 0.9656 | 0.9852 | 0.7933 | 0.4798 |
| AlphaFold-FineTune | 26216 | 2202 | 13280 | 12150 | 0.7335 | 0.6833 | 0.8578 | 0.9225 | 0.7705 | 0.9157 | 0.7851 | 0.4222 |
| STRUMP-I | 35149 | 2056 | 13426 | 3217 | 0.9021 | 0.9161 | 0.8672 | 0.9447 | 0.9604 | 0.9836 | 0.9302 | 0.7555 |
| MHCFlurry - NetMHCpan | 28703 | 783 | 14699 | 9663 | 0.806 | 0.7481 | 0.9734 | 0.9494 | 0.9132 | 0.9678 | 0.8460 | 0.5561 |
| MHCFlurry - HLAthena | 33950 | 1249 | 14233 | 4416 | 0.8948 | 0.8849 | 0.9193 | 0.9645 | 0.9489 | 0.9808 | 0.9230 | 0.7328 |
| NetMHCpan HLAthena | 27203 | 414 | 15068 | 11163 | 0.785 | 0.709 | 0.9733 | 0.985 | 0.8679 | 0.9586 | 0.8245 | 0.5242 |
| STRUMP-I - HLAthena | 32977 | 502 | 14980 | 5389 | 0.8906 | 0.8595 | 0.9676 | 0.985 | 0.947 | 0.9828 | 0.9180 | 0.7211 |
| STRUMP-I - MHCFlurry | 33509 | 1086 | 14396 | 4859 | 0.8896 | 0.8734 | 0.9299 | 0.9686 | 0.9488 | 0.9814 | 0.9185 | 0.7202 |
| STRUMP-I - NetMHCpan | 26921 | 386 | 15096 | 11445 | 0.7803 | 0.7017 | 0.9751 | 0.9859 | 0.8678 | 0.9588 | 0.8198 | 0.5169 |
| MHCFlurry - NetMHCpan - HLAthena | 26954 | 414 | 15068 | 11412 | 0.7804 | 0.7025 | 0.9733 | 0.9849 | 0.8632 | 0.9573 | 0.8201 | 0.5167 |
| STRUMP-I - MHCFlurry - NetMHCpan | 26572 | 371 | 15111 | 11794 | 0.7741 | 0.6926 | 0.976 | 0.9862 | 0.8631 | 0.9575 | 0.8137 | 0.5072 |
| STRUMP-I - MHCFlurry - HLAthena | 31904 | 469 | 15013 | 6462 | 0.8713 | 0.8316 | 0.9697 | 0.9855 | 0.932 | 0.9783 | 0.9020 | 0.6803 |
| STRUMP-I - NetMHCpan - HLAthena | 25553 | 213 | 15269 | 12813 | 0.7581 | 0.666 | 0.9862 | 0.9917 | 0.8451 | 0.9535 | 0.7969 | 0.4843 |
| STRUMP-I - MHCFlurry - NetMHCpan - HLAthena | 25314 | 213 | 15269 | 13052 | 0.7537 | 0.6598 | 0.9862 | 0.9917 | 0.8451 | 0.9535 | 0.7924 | 0.4777 |

Table S3. Performance metrics on the validation set from the training data

| Model | TP | FP | TN | FN | Accuracy | Recall | Specificity | Precision | AUC | PRAUC | F1 | MCC |
|---|---|---|---|---|---|---|---|---|---|---|---|---|
| NetMHCpan RankEL | 23945 | 545 | 11876 | 6747 | 0.8309 | 0.7802 | 0.9561 | 0.9777 | 0.9479 | 0.9745 | 0.8679 | 0.6013 |
| NetMHCpan Affinity | 23101 | 664 | 11757 | 7591 | 0.8085 | 0.7527 | 0.9465 | 0.9721 | 0.9395 | 0.9745 | 0.8484 | 0.5602 |
| MHCFlurry Affinity | 28345 | 1429 | 10922 | 2347 | 0.9124 | 0.9235 | 0.885 | 0.952 | 0.962 | 0.9836 | 0.9376 | 0.7794 |
| MHCFlurry Affinity Percentile | 28995 | 3120 | 9301 | 1500 | 0.8923 | 0.9508 | 0.7488 | 0.9028 | 0.9507 | 0.9793 | 0.9262 | 0.7503 |
| HLAthena MSi | 28027 | 1164 | 11257 | 2665 | 0.9112 | 0.9132 | 0.9063 | 0.9601 | 0.9686 | 0.987 | 0.9361 | 0.7740 |
| HLAthena prank MSi | 20330 | 127 | 12294 | 10362 | 0.7567 | 0.6624 | 0.9898 | 0.9938 | 0.9665 | 0.9859 | 0.7949 | 0.4829 |
| AlphaFold-FineTune | 28183 | 1732 | 10689 | 2509 | 0.9016 | 0.9183 | 0.8841 | 0.952 | 0.9685 | 0.9867 | 0.9300 | 0.7555 |
| STRUMP-I | 22787 | 610 | 11811 | 7905 | 0.8025 | 0.7424 | 0.9509 | 0.9739 | 0.913 | 0.9678 | 0.8426 | 0.5503 |
| MHCFlurry - NetMHCpan | 27098 | 1009 | 11412 | 3594 | 0.8932 | 0.8829 | 0.9188 | 0.9641 | 0.9494 | 0.9816 | 0.9217 | 0.7294 |
| MHCFlurry - HLAthena | 2155 | 318 | 12103 | 9137 | 0.7807 | 0.7023 | 0.9744 | 0.9855 | 0.8657 | 0.9583 | 0.3131 | 0.1264 |
| NetMHCpan HLAthena | 26409 | 437 | 11984 | 4283 | 0.8905 | 0.8605 | 0.9648 | 0.9837 | 0.946 | 0.9824 | 0.9180 | 0.7211 |
| STRUMP-I - HLAthena | 26841 | 919 | 11502 | 3851 | 0.8894 | 0.8745 | 0.926 | 0.9669 | 0.9488 | 0.9815 | 0.9184 | 0.7201 |
| STRUMP-I - MHCFlurry | 21439 | 317 | 12104 | 9253 | 0.778 | 0.6985 | 0.9745 | 0.9854 | 0.865 | 0.9579 | 0.8175 | 0.5132 |
| STRUMP-I - NetMHCpan | 21359 | 313 | 12108 | 9333 | 0.7763 | 0.6959 | 0.9748 | 0.9856 | 0.8611 | 0.957 | 0.8158 | 0.5105 |
| MHCFlurry - NetMHCpan - HLAthena | 21172 | 300 | 12121 | 9520 | 0.7722 | 0.6898 | 0.9758 | 0.986 | 0.8605 | 0.9566 | 0.8117 | 0.5043 |
| STRUMP-I - MHCFlurry - NetMHCpan | 25549 | 399 | 12022 | 5143 | 0.8715 | 0.8324 | 0.9679 | 0.9846 | 0.931 | 0.9781 | 0.9022 | 0.6808 |
| STRUMP-I - MHCFlurry - HLAthena | 20335 | 176 | 12245 | 10357 | 0.7557 | 0.6626 | 0.9858 | 0.9914 | 0.8452 | 0.9535 | 0.7943 | 0.4807 |
| STRUMP-I - NetMHCpan - HLAthena | 20152 | 174 | 12247 | 10540 | 0.7515 | 0.6566 | 0.986 | 0.9914 | 0.842 | 0.9525 | 0.7900 | 0.4744 |
| STRUMP-I - MHCFlurry - NetMHCpan - HLAthena | 23945 | 545 | 11876 | 6747 | 0.8309 | 0.7802 | 0.9561 | 0.9777 | 0.9479 | 0.9745 | 0.8679 | 0.6013 |

Table S4. Allele representation in the PRIME and TESLA datasets

| Allele | Allele training size | Positive ratio | PRIME | TESLA | In-House |
|---|---|---|---|---|---|
| A0101: | 2811 | 0.378571429 | X | X | |
| A0103: | 0 | -- | X | | |
| A0201: | 7788 | 0.529872674 | X | X | X |
| A0202: | 3356 | 0.749928795 | X | | |
| A0203: | 3935 | 0.668073486 | X | | |
| A0206: | 3449 | 0.66731789 | X | | X |
| A0211: | 1522 | 0.779153094 | X | | |
| A0301: | 4578 | 0.466129384 | X | X | |
| A1101: | 5148 | 0.619538256 | X | X | X |
| A2301: | 2370 | 0.703339041 | X | | |
| A2402: | 2638 | 0.624148372 | X | X | X |
| A2501: | 1135 | 0.54787234 | X | | |
| A2601: | 2691 | 0.380395026 | X | | |
| A2902: | 1704 | 0.519666269 | X | | |
| A3001: | 1626 | 0.676962677 | X | | |
| A3002: | 2064 | 0.765936255 | X | | |
| A3101: | 3882 | 0.414314809 | X | | X |
| A3201: | 1621 | 0.830759728 | X | | |
| A3301: | 3097 | 0.526565144 | X | | |
| A6601: | 1050 | 0.969387755 | X | X | |
| A6801: | 2757 | 0.650817236 | X | | |
| A6802: | 3412 | 0.455362156 | X | | |
| A6812: | 0 | -- | X | | |
| A6901: | 1462 | 0.104579631 | X | | |
| B0702: | 2969 | 0.531564809 | | | X |
| B0801: | 1747 | 0.446418338 | X | X | |
| B1402: | 904 | 0.95049505 | X | | |
| B1501: | 4142 | 0.65631769 | X | | X |
| B1502: | 914 | 0.974054054 | X | | |
| B1517: | 1797 | 0.654980523 | X | | |
| B1801: | 2514 | 0.525834658 | X | | |
| B2702: | 2 | 0 | X | | |
| B2705: | 2104 | 0.51161688 | X | X | |
| B3501: | 1385 | 0.6534867 | X | | |
| B3503: | 781 | 0.996168582 | X | | |
| B3701: | 900 | 0.986827662 | X | | |
| B0702: | 2969 | 0.531564809 | X | | |
| B3801: | 1386 | 0.940373563 | X | | |
| B3901: | 806 | 0.175308642 | X | | |

| | | | | | |
|---|---|---|---|---|---|
| B3906: | 22 | 0.772727273 | X | | X |
| B4001: | 3328 | 0.636744464 | X | | |
| B4002: | 2526 | 0.92724397 | X | | |
| B4101: | 0 | -- | X | | |
| B4102: | 0 | -- | X | | |
| B4402: | 1647 | 0.513085819 | X | X | X |
| B4403: | 1098 | 0.687155963 | X | | |
| B4501: | 1047 | 0.832221163 | X | | |
| B4901: | 2219 | 1 | X | | X |
| B5101: | 2282 | 0.485989492 | X | | |
| B5301: | 1841 | 0.788648649 | X | | |
| B5601: | 1003 | 1 | X | | |
| B5701: | 2031 | 0.418799213 | X | X | |
| B5801: | 2524 | 0.557364033 | X | | |
| C0102: | 768 | 1 | X | | |
| C0202: | 550 | 1 | X | | |
| C0303: | 1302 | 0.982496195 | X | | |
| C0304: | 1438 | 1 | X | | |
| C0401: | 1410 | 0.786978061 | X | | |
| C0501: | 915 | 0.923747277 | X | | |
| C0602: | 943 | 0.925498426 | X | | |
| C0701: | 430 | 1 | X | | |
| C0702: | 660 | 0.947049924 | X | | |
| C0802: | 1836 | 0.98212351 | X | | |
| C1203: | 1395 | 0.98782235 | X | | |
| C1402: | 960 | 0.973305955 | X | | |
| C1403: | 1545 | 1 | X | | |
| C1502: | 2112 | 0.959810875 | X | | |
| C1601: | 1755 | 1 | X | | |

Table S5. Perfomance metrics on the test set from average represented alleles in the training data (<2000 pMHCs represented)

| Model | TP | FP | TN | FN | Accuracy | Recall | Specificity | Precision | AUC | PRAUC | F1 | MCC |
|---|---|---|---|---|---|---|---|---|---|---|---|---|
| NetMHCpan RankEL | 17388 | 293 | 4585 | 3788 | 0.8434 | 0.8211 | 0.9399 | 0.9834 | 0.9466 | 0.9866 | 0.8950 | 0.5809 |
| NetMHCpan Affinity | 15377 | 185 | 4693 | 5799 | 0.7703 | 0.7262 | 0.9621 | 0.9811 | 0.941 | 0.9853 | 0.8371 | 0.4693 |
| MHCFlurry Affinity | 19866 | 525 | 4353 | 1310 | 0.9296 | 0.9381 | 0.8924 | 0.9743 | 0.9714 | 0.9929 | 0.9448 | 0.7708 |
| MHCFlurry Affinity Percentile | 19999 | 1418 | 3460 | 918 | 0.9094 | 0.9561 | 0.7093 | 0.9338 | 0.9477 | 0.9863 | 0.9559 | 0.7024 |
| HLAthena MSi | 20303 | 523 | 4355 | 873 | 0.9464 | 0.9588 | 0.8928 | 0.9749 | 0.9785 | 0.9944 | 0.9668 | 0.8226 |
| HLAthena prank MSi | 15603 | 70 | 4808 | 5573 | 0.7834 | 0.7368 | 0.9856 | 0.9955 | 0.9765 | 0.9937 | 0.8469 | 0.4952 |
| AlphaFold-FineTune | 14237 | 707 | 4171 | 6939 | 0.7065 | 0.6723 | 0.8551 | 0.9527 | 0.8342 | 0.9492 | 0.7883 | 0.3495 |
| STRUMP-I | 20273 | 443 | 4435 | 903 | 0.9413 | 0.9578 | 0.8671 | 0.9701 | 0.9785 | 0.9949 | 0.9679 | 0.8284 |

Table S6. Performance metrics on the test set from very lowly represented alleles in the training data (<500 pMHCs represented)

| Model | TP | FP | TN | FN | Accuracy | Recall | Specificity | Precision | AUC | PRAUC | F1 | MCC |
|---|---|---|---|---|---|---|---|---|---|---|---|---|
| NetMHCpan RankEL | 315 | 15 | 617 | 374 | 0.7055 | 0.4572 | 0.9763 | 0.9545 | 0.8342 | 0.8706 | 0.6183 | 0.3461 |
| NetMHCpan Affinity | 630 | 248 | 384 | 59 | 0.7676 | 0.9144 | 0.6076 | 0.7175 | 0.8719 | 0.8776 | 0.8041 | 0.6234 |
| MHCFlurry Affinity | 602 | 54 | 578 | 87 | 0.8933 | 0.8737 | 0.9146 | 0.9177 | 0.9507 | 0.9518 | 0.8952 | 0.7685 |
| MHCFlurry Affinity Percentile | 630 | 248 | 384 | 59 | 0.7676 | 0.9144 | 0.6076 | 0.7175 | 0.8719 | 0.8776 | 0.8041 | 0.6234 |
| HLAthena MSi | 603 | 93 | 539 | 86 | 0.8645 | 0.8752 | 0.8528 | 0.8664 | 0.9294 | 0.932 | 0.8708 | 0.7321 |
| HLAthena prank MSi | 425 | 17 | 615 | 264 | 0.7873 | 0.6168 | 0.9731 | 0.9615 | 0.9225 | 0.9326 | 0.7515 | 0.5002 |
| AlphaFold-FineTune | 363 | 131 | 501 | 326 | 0.654 | 0.5269 | 0.7927 | 0.7348 | 0.7094 | 0.9335 | 0.6137 | 0.2794 |
| STRUMP-I | 641 | 18 | 614 | 48 | 0.95 | 0.9303 | 0.9715 | 0.9727 | 0.9847 | 0.9891 | 0.9510 | 0.8812 |

Table S7. Performance metrics on the test set from overrepresented alleles in the  training data (>=2000) pMHCs represented)

| Model | TP | FP | TN | FN | Accuracy | Recall | Specificity | Precision | AUC | PRAUC | F1 | MCC |
|---|---|---|---|---|---|---|---|---|---|---|---|---|
| NetMHCpan RankEL | 12597 | 387 | 10217 | 4593 | 0.8208 | 0.7328 | 0.9635 | 0.9702 | 0.9439 | 0.9662 | 0.8350 | 0.5892 |
| NetMHCpan Affinity | 13718 | 648 | 9956 | 3472 | 0.8518 | 0.798 | 0.9389 | 0.9549 | 0.9483 | 0.9676 | 0.8694 | 0.6549 |
| MHCFlurry Affinity | 15614 | 1285 | 9319 | 1576 | 0.8971 | 0.9083 | 0.8788 | 0.924 | 0.9527 | 0.9685 | 0.9161 | 0.7766 |
| MHCFlurry Affinity Percentile | 16276 | 2510 | 8094 | 914 | 0.8768 | 0.9468 | 0.7633 | 0.8664 | 0.9483 | 0.9675 | 0.9048 | 0.7705 |
| HLAthena MSi | 14800 | 906 | 9698 | 2390 | 0.8814 | 0.861 | 0.9146 | 0.9423 | 0.9538 | 0.9713 | 0.8998 | 0.7264 |
| HLAthena prank MSi | 9736 | 106 | 10498 | 7454 | 0.728 | 0.5664 | 0.99 | 0.9892 | 0.9512 | 0.9694 | 0.7203 | 0.4276 |
| AlphaFold-FineTune | 11979 | 1495 | 9109 | 5211 | 0.7587 | 0.6969 | 0.859 | 0.889 | 0.846 | 0.8904 | 0.7813 | 0.4783 |
| STRUMP-I | 14567 | 1040 | 9564 | 2623 | 0.8682 | 0.8474 | 0.9019 | 0.9334 | 0.9447 | 0.9628 | 0.8883 | 0.6990 |

Table S8. Perfomance metrics on the TESLA peptide set

| Model | TP | FP | TN | FN | Accuracy | Recall | Specificity | Precision | AUC | PRAUC | F1 | MCC |
|---|---|---|---|---|---|---|---|---|---|---|---|---|
| NetMHCpan RankEL | 245 | 35 | 99 | 121 | 0.688 | 0.6694 | 0.7388 | 0.875 | 0.7519 | 0.8809 | 0.7585 | 0.318581 |
| NetMHCpan Affinity | 359 | 84 | 50 | 7 | 0.818 | 0.9809 | 0.3731 | 0.8104 | 0.8492 | 0.9371 | 0.8875 | 0.542786 |
| MHCFlurry Affinity | 332 | 66 | 68 | 34 | 0.8 | 0.9071 | 0.5075 | 0.8342 | 0.8272 | 0.9292 | 0.8691 | 0.475167 |
| MHCFlurry Affinity Percentile | 350 | 94 | 40 | 16 | 0.78 | 0.9563 | 0.2985 | 0.7883 | 0.801 | 0.9094 | 0.8642 | 0.394134 |
| HLAthena MSi | 269 | 74 | 60 | 97 | 0.658 | 0.735 | 0.4478 | 0.7843 | 0.6686 | 0.838 | 0.7588 | 0.168819 |
| HLAthena prank MSi | 99 | 14 | 120 | 267 | 0.438 | 0.2705 | 0.8955 | 0.8761 | 0.6671 | 0.8377 | 0.4134 | 0.097688 |
| AlphaFold-FineTune | 254 | 49 | 85 | 112 | 0.678 | 0.694 | 0.6343 | 0.8383 | 0.6911 | 0.8312 | 0.7593 | 0.270778 |
| STRUMP-I | 296 | 82 | 52 | 70 | 0.696 | 0.8087 | 0.3881 | 0.7831 | 0.5894 | 0.8659 | 0.7957 | 0.206255 |
| MHCFlurry - NetMHCpan | 346 | 61 | 73 | 20 | 0.838 | 0.9454 | 0.5448 | 0.8501 | 0.8759 | 0.9519 | 0.8453 | 0.588333 |
| MHCFlurry - HLAthena | 265 | 58 | 76 | 101 | 0.682 | 0.724 | 0.5672 | 0.7204 | 0.7322 | 0.867 | 0.7692 | 0.253378 |
| NetMHCpan HLAthena | 266 | 51 | 83 | 100 | 0.698 | 0.7268 | 0.6194 | 0.8391 | 0.7625 | 0.8744 | 0.7789 | 0.296229 |
| STRUMP-I - HLAthena | 234 | 53 | 81 | 132 | 0.63 | 0.6393 | 0.6045 | 0.8153 | 0.6219 | 0.8593 | 0.7167 | 0.193391 |
| STRUMP-I - MHCFlurry | 287 | 63 | 71 | 79 | 0.716 | 0.7842 | 0.5299 | 0.82 | 0.657 | 0.8811 | 0.8017 | 0.296785 |
| STRUMP-I - NetMHCpan | 292 | 57 | 77 | 74 | 0.738 | 0.7978 | 0.5746 | 0.8367 | 0.6862 | 0.8912 | 0.8168 | 0.35085 |
| MHCFlurry - NetMHCpan - HLAthena | 262 | 43 | 91 | 104 | 0.706 | 0.7158 | 0.6791 | 0.859 | 0.7935 | 0.8905 | 0.7809 | 0.3274 |
| STRUMP-I - MHCFlurry - NetMHCpan | 284 | 42 | 92 | 82 | 0.752 | 0.776 | 0.6866 | 0.8712 | 0.7313 | 0.9056 | 0.8208 | 0.405893 |
| STRUMP-I - MHCFlurry - HLAthena | 230 | 44 | 90 | 136 | 0.64 | 0.6284 | 0.6716 | 0.8394 | 0.65 | 0.8699 | 0.7186 | 0.231048 |
| STRUMP-I - NetMHCpan - HLAthena | 231 | 38 | 96 | 135 | 0.654 | 0.6311 | 0.7164 | 0.8587 | 0.6738 | 0.8799 | 0.7276 | 0.264718 |
| STRUMP-I - MHCFlurry - NetMHCpan - HLAthena | 227 | 31 | 103 | 139 | 0.66 | 0.6202 | 0.7687 | 0.8798 | 0.6944 | 0.889 | 0.7276 | 0.289371 |

Table S9. Performance metrics on the PRIME peptide set

| Model | TP | FP | TN | FN | Accuracy | Recall | Specificity | Precision | AUC | PRAUC | F1 | MCC |
|---|---|---|---|---|---|---|---|---|---|---|---|---|
| NetMHCpan RankEL | 2330 | 38 | 2749 | 2573 | 0.6605 | 0.4752 | 0.9864 | 0.984 | 0.9639 | 0.974 | 0.6415 | 0.334028 |
| NetMHCpan Affinity | 3688 | 45 | 2742 | 1215 | 0.8362 | 0.7522 | 0.9839 | 0.9879 | 0.9838 | 0.9884 | 0.8541 | 0.617718 |
| MHCFlurry Affinity | 3849 | 109 | 2678 | 1054 | 0.8488 | 0.785 | 0.9609 | 0.9725 | 0.9609 | 0.9754 | 0.8688 | 0.644599 |
| MHCFlurry Affinity Percentile | 4273 | 190 | 2597 | 630 | 0.8934 | 0.8715 | 0.9318 | 0.9574 | 0.9593 | 0.9723 | 0.9124 | 0.74656 |
| HLAthena MSi | 3202 | 52 | 2732 | 1701 | 0.772 | 0.6531 | 0.9813 | 0.984 | 0.9759 | 0.982 | 0.7851 | 0.50274 |
| HLAthena prank MSi | 860 | 8 | 2779 | 4043 | 0.4732 | 0.1754 | 0.9971 | 0.9908 | 0.9743 | 0.9804 | 0.298 | 0.110277 |
| AlphaFold-FineTune | 3315 | 80 | 2707 | 1588 | 0.7831 | 0.6761 | 0.9713 | 0.9764 | 0.9196 | 0.9532 | 0.7989 | 0.521516 |
| STRUMP-I | 3941 | 1230 | 1557 | 962 | 0.715 | 0.8038 | 0.5587 | 0.7621 | 0.6812 | 0.8455 | 0.7824 | 0.381253 |
| MHCFlurry - NetMHCpan | 3556 | 41 | 2746 | 1347 | 0.8195 | 0.7253 | 0.9853 | 0.9886 | 0.9207 | 0.9633 | 0.7787 | 0.586338 |
| MHCFlurry - HLAthena | 3076 | 46 | 2741 | 1827 | 0.7564 | 0.6274 | 0.9835 | 0.9853 | 0.9205 | 0.9633 | 0.7666 | 0.477146 |
| NetMHCpan HLAthena | 2753 | 25 | 2762 | 2150 | 0.7126 | 0.5542 | 0.9914 | 0.9912 | 0.857 | 0.9439 | 0.7168 | 0.416188 |
| STRUMP-I - HLAthena | 2729 | 43 | 2744 | 2174 | 0.7117 | 0.5566 | 0.9846 | 0.9845 | 0.7706 | 0.9119 | 0.7111 | 0.407387 |
| STRUMP-I - MHCFlurry | 3470 | 138 | 2649 | 1433 | 0.7957 | 0.7077 | 0.9505 | 0.9618 | 0.8291 | 0.9279 | 0.8154 | 0.543875 |
| STRUMP-I - NetMHCpan | 3081 | 41 | 2746 | 1822 | 0.7577 | 0.6284 | 0.9853 | 0.9869 | 0.8068 | 0.9261 | 0.7679 | 0.479344 |
| MHCFlurry - NetMHCpan - HLAthena | 2717 | 24 | 2763 | 2186 | 0.7126 | 0.5542 | 0.9914 | 0.9912 | 0.857 | 0.9439 | 0.7109 | 0.409389 |
| STRUMP-I - MHCFlurry - NetMHCpan | 2983 | 38 | 2749 | 1920 | 0.7454 | 0.6084 | 0.9864 | 0.9874 | 0.7974 | 0.9227 | 0.7529 | 0.45952 |
| STRUMP-I - MHCFlurry - HLAthena | 2632 | 37 | 2750 | 2271 | 0.6999 | 0.5368 | 0.9867 | 0.9861 | 0.7618 | 0.9091 | 0.6952 | 0.390051 |
| STRUMP-I - NetMHCpan - HLAthena | 2390 | 24 | 3763 | 2513 | 0.6701 | 0.4875 | 0.9914 | 0.9901 | 0.7394 | 0.9022 | 0.6533 | 0.373731 |
| STRUMP-I - MHCFlurry - NetMHCpan - HLAthena | 2362 | 23 | 2764 | 2541 | 0.6666 | 0.4817 | 0.9917 | 0.9904 | 0.7367 | 0.9013 | 0.6482 | 0.343194 |

Table S10 VACINUS cancer neoantigen dataset

| allele | peptide | Kd | Binder |
|---|---|---|---|
| A0201 | FLPDPLIFQM | 4.3 | TRUE |
| A0201 | KLSPEEVEYL | 5.8 | TRUE |
| A0201 | MLGQVKYNL | 0.55 | TRUE |
| A0201 | SLSQLLHEM | 1.9 | TRUE |
| A0201 | SLVTFQEPL | 0.1 | TRUE |
| A0201 | SMLGQVKYNL | 5.8 | TRUE |
| A0201 | control | 0.9 | TRUE |
| A0206 | GVLKFACLV | 1.4 | TRUE |
| A0206 | HVPDYLVPSA | 1.55 | TRUE |
| A0206 | LQMTSYHFA | 0.8 | TRUE |
| A0206 | YLVPSALRGL | 0.4 | TRUE |
| A0206 | YTLDHFSKSL | 1.65 | TRUE |
| A0206 | control | 0.8 | TRUE |
| A1101 | AIFGWGHNK | 0.1 | TRUE |
| A1101 | ASADLPPPK | 0.95 | TRUE |
| A1101 | ATTSLAVATR | 50000 | FALSE |
| A1101 | GAIFGWGHNK | 2.3 | TRUE |
| A1101 | GVFRAHLFR | 0.8 | TRUE |
| A1101 | KTKPLPVLGK | 2.65 | TRUE |
| A1101 | LLCFWNSYAK | 50000 | FALSE |
| A1101 | LSISGFLSK | 0.55 | TRUE |
| A1101 | NSYAKTLASK | 50000 | FALSE |
| A1101 | QIANIDHITK | 1.55 | TRUE |
| A1101 | RAFSTAPWR | 1000 | FALSE |
| A1101 | RSSGLWPFTR | 1000 | FALSE |
| A1101 | RSWFPSAPR | 50000 | FALSE |
| A1101 | RTSRLTSLR | 2.1 | TRUE |
| A1101 | RVPAFAFVR | 0.25 | TRUE |
| A1101 | RVSNIFPKQK | 3 | TRUE |
| A1101 | SAFAALHAK | 0.1 | TRUE |
| A1101 | SAMTTSSSQK | 0.3 | TRUE |
| A1101 | SIPTRVINHK | 0.55 | TRUE |
| A1101 | SVNSIVLLFK | 87.85 | TRUE |
| A1101 | TTSLAVATR | 50000 | FALSE |
| A1101 | VLVWHTWTEK | 50000 | FALSE |
| A1101 | control | 2.6 | TRUE |
| A2402 | AYLGRTLLV | 15.45 | TRUE |
| A2402 | FYINVERVEW | 22.75 | TRUE |
| A2402 | GWFSFFMPF | 50000 | FALSE |
| A2402 | HWFRKGLRF | 50000 | FALSE |

| | | | |
|---|---|---|---|
| A2402 | IAPGWFSFF | 50000 | FALSE |
| A2402 | IFAQEALALF | 0.25 | TRUE |
| A2402 | KWLDLTYSF | 55.25 | TRUE |
| A2402 | KYAMAVGSL | 0.15 | TRUE |
| A2402 | KYPVGVHVL | 50000 | FALSE |
| A2402 | LYFSSNMREF | 0.15 | TRUE |
| A2402 | QWLFVSLNMF | 50000 | FALSE |
| A2402 | RWPEACSIW | 50000 | FALSE |
| A2402 | SYMPKMAVI | 0.6 | TRUE |
| A2402 | TYDHRAYSSF | 50000 | FALSE |
| A2402 | TYQGFISQW | 0.1 | TRUE |
| A2402 | VFPHYSSKF | 0.1 | TRUE |
| A2402 | control | 40.225 | TRUE |
| A3101 | AFVTRSFPR | 0.65 | TRUE |
| A3101 | AMLKAPHPR | 1.05 | TRUE |
| A3101 | AQFPGVCGLR | 3.5 | TRUE |
| A3101 | ATWRRCTPR | 2.05 | TRUE |
| A3101 | GFFGPMKYYR | 1.15 | TRUE |
| A3101 | KLKDLIWFR | 1.2 | TRUE |
| A3101 | KMHSGEFAR | 3.3 | TRUE |
| A3101 | KSHMGRLLAR | 69.9 | TRUE |
| A3101 | KSQEKLLRAR | 7.95 | TRUE |
| A3101 | KTIGFDFAK | 1.05 | TRUE |
| A3101 | KVSNITHSR | 3.9 | TRUE |
| A3101 | KYGKDIAAYR | 1.8 | TRUE |
| A3101 | NTRTPCTPR | 8.6 | TRUE |
| A3101 | RAPRSPFLR | 50000 | FALSE |
| A3101 | RLKALRRTR | 4.5 | TRUE |
| A3101 | RLNTREFHTR | 1.3 | TRUE |
| A3101 | RSMSANPKQR | 16.2 | TRUE |
| A3101 | RSRGQLLRR | 3.45 | TRUE |
| A3101 | RSTPAQTAR | 9.6 | TRUE |
| A3101 | RSWTSKGWR | 55.3 | TRUE |
| A3101 | RSWTSKGWRR | 2.1 | TRUE |
| A3101 | SVKPIYKKR | 1.4 | TRUE |
| A3101 | TIRENYIHR | 8.75 | TRUE |
| A3101 | VAFVTRSFPR | 56.05 | TRUE |
| A3101 | VTRSFPRPR | 5.5 | TRUE |
| A3101 | control | 13.53333 | TRUE |
| B1501 | CQARHARFF | 0.8 | TRUE |
| B1501 | GQIGVHTPL | 0.1 | TRUE |

| | | | |
|---|---|---|---|
| B1501 | KSNGTSVVF | 0.1 | TRUE |
| B1501 | LQKNAAAAV | 0.1 | TRUE |
| B1501 | NLKDFISCF | 28.05 | TRUE |
| B1501 | RLIHGEVPM | 0.4 | TRUE |
| B1501 | RLYQDTSLM | 0.65 | TRUE |
| B1501 | RMIVLPHSL | 2.8 | TRUE |
| B1501 | RQRSSGLWPF | 1.45 | TRUE |
| B1501 | SLGAHISEF | 5.45 | TRUE |
| B1501 | SLISAPRSM | 0.2 | TRUE |
| B1501 | SLLCFWNSY | 270.9 | TRUE |
| B1501 | SQLRYARTM | 0.2 | TRUE |
| B1501 | SQMQMTLAY | 3.575 | TRUE |
| B1501 | SQREEYRGF | 2.8 | TRUE |
| B1501 | SVNSIVLLF | 50000 | FALSE |
| B1501 | TQAPAFLEL | 2.1 | TRUE |
| B1501 | VAMYGGYAGY | 2 | TRUE |
| B1501 | VVFNPSVLHY | 0.4 | TRUE |
| B1501 | control | 1.225 | TRUE |
| B4001 | AETPGAPPL | 0.15 | TRUE |
| B4001 | GEATVHEPV | 0.35 | TRUE |
| B4001 | GEATVHEPVL | 0.3 | TRUE |
| B4001 | GEFTSTRAVL | 0.35 | TRUE |
| B4001 | IETEKTLTL | 0.45 | TRUE |
| B4001 | IETNCIYHL | 0.6 | TRUE |
| B4001 | KEATSNQDLL | 0.35 | TRUE |
| B4001 | KEILWPLPI | 0.15 | TRUE |
| B4001 | KETQWLFVSL | 1.45 | TRUE |
| B4001 | LEAQVEAAL | 3.1 | TRUE |
| B4001 | MESECGVVI | 0.05 | TRUE |
| B4001 | QESPEVLLTL | 0.2 | TRUE |
| B4001 | REAFCRQNL | 0.3 | TRUE |
| B4001 | REDEALLLVV | 0.8 | TRUE |
| B4001 | REHLSEEVIL | 0.2 | TRUE |
| B4001 | SEFACNAAL | 0.05 | TRUE |
| B4001 | SELAQINSL | 5.85 | TRUE |
| B4001 | WEASVVTFF | 6.15 | TRUE |
| B4001 | control | 19.6 | TRUE |
| B4403 | REAPTMTLF | 6.8 | TRUE |
| B4403 | SENLAKRLY | 127.15 | TRUE |
| B4403 | WENPFYYLF | 7.9 | TRUE |
| B4403 | control | 589.45 | FALSE |

| | | | |
|---|---|---|---|
| B5101 | CPFFMSVNI | 214.4 | TRUE |
| B5101 | CPIFQISNI | 50000 | FALSE |
| B5101 | FPAPDTHNI | 22.1 | TRUE |
| B5101 | IPHEMLMGI | 58.65 | TRUE |
| B5101 | LPMVCTASV | 24.8 | TRUE |
| B5101 | LPTENELTI | 16.75 | TRUE |
| B5101 | control | 28.8 | TRUE |

Table S11. Performance metrics on the VACINUS peptide set

| Model | TP | FP | TN | FN | Accuracy | Recall | Specificity | Precision | AUC | PRAUC | F1 | MCC |
|---|---|---|---|---|---|---|---|---|---|---|---|---|
| NetMHCpan RankEL | 170 | 20 | 16 | 34 | 0.775 | 0.8333 | 0.4444 | 0.8947 | 0.7225 | 0.9369 | 0.8629 | 0.235702 |
| NetMHCpan Affinity | 204 | 36 | 0 | 0 | 0.85 | 1 | 0 | 0.85 | 0.7952 | 0.96 | 0.9189 | -- |
| MHCFlurry Affinity | 204 | 36 | 0 | 0 | 0.85 | 1 | 0 | 0.85 | 0.6852 | 0.9282 | 0.9189 | -- |
| MHCFlurry Affinity Percentile | 204 | 36 | 0 | 0 | 0.85 | 1 | 0 | 0.85 | 0.6759 | 0.9316 | 0.9189 | -- |
| HLAthena MSi | 184 | 26 | 10 | 20 | 0.8083 | 0.902 | 0.2778 | 0.8762 | 0.6748 | 0.9019 | 0.8889 | 0.196894 |
| HLAthena prank MSi | 64 | 6 | 30 | 140 | 0.3917 | 0.3137 | 0.8333 | 0.9143 | 0.6825 | 0.9067 | 0.4672 | 0.067673 |
| AlphaFold-FineTune | 166 | 26 | 10 | 38 | 0.7333 | 0.8137 | 0.2778 | 0.8646 | 0.6008 | 0.9052 | 0.8384 | 0.079244 |
| STRUMP-I | 186 | 28 | 8 | 18 | 0.8083 | 0.9118 | 0.2222 | 0.8692 | 0.567 | 0.928 | 0.89 | 0.157662 |
| MHCFlurry - NetMHCpan | 204 | 36 | 0 | 0 | 0.85 | 1 | 0 | 0.85 | 0.7952 | 0.9018 | 0.9189 | -- |
| MHCFlurry - HLAthena | 184 | 26 | 10 | 20 | 0.8083 | 0.902 | 0.2778 | 0.8762 | 0.6748 | 0.9018 | 0.8889 | 0.196894 |
| NetMHCpan HLAthena | 184 | 26 | 10 | 20 | 0.8083 | 0.902 | 0.2778 | 0.8762 | 0.6748 | 0.9018 | 0.8889 | 0.196894 |
| STRUMP-I - HLAthena | 170 | 20 | 16 | 34 | 0.775 | 0.8333 | 0.4444 | 0.8947 | 0.6389 | 0.9349 | 0.8629 | 0.235702 |
| STRUMP-I - MHCFlurry | 186 | 28 | 8 | 18 | 0.8083 | 0.9118 | 0.2222 | 0.8692 | 0.567 | 0.928 | 0.89 | 0.157662 |
| STRUMP-I - NetMHCpan | 186 | 28 | 8 | 18 | 0.8083 | 0.9118 | 0.2222 | 0.8692 | 0.567 | 0.928 | 0.89 | 0.157662 |
| MHCFlurry - NetMHCpan - HLAthena | 184 | 26 | 10 | 20 | 0.8083 | 0.902 | 0.2778 | 0.8762 | 0.6748 | 0.9018 | 0.8889 | 0.196894 |
| STRUMP-I - MHCFlurry - NetMHCpan | 186 | 28 | 8 | 18 | 0.8083 | 0.9118 | 0.2222 | 0.8692 | 0.567 | 0.928 | 0.89 | 0.157662 |
| STRUMP-I - MHCFlurry - HLAthena | 170 | 20 | 16 | 34 | 0.775 | 0.8333 | 0.4444 | 0.8947 | 0.6389 | 0.9349 | 0.863 | 0.235702 |
| STRUMP-I - NetMHCpan - HLAthena | 170 | 20 | 16 | 34 | 0.775 | 0.8333 | 0.4444 | 0.8947 | 0.6389 | 0.9349 | 0.863 | 0.235702 |
| STRUMP-I - MHCFlurry - NetMHCpan - HLAthena | 170 | 20 | 16 | 34 | 0.775 | 0.8333 | 0.4444 | 0.8947 | 0.6389 | 0.9349 | 0.863 | 0.235702 |

Table S12. LightGBM hyperparameters used for model training

| Hyperparameter | Binding prediction value |
| --- | --- |
| boosting_type | dart |
| lambda_l1 | 0.002185954 |
| lambda_l2 | 1.18E-05 |
| num_leaves | 256 |
| feature_fraction | 0.646750402 |
| bagging_fraction | 0.933824118 |
| bagging_freq | 6 |
| min_data_in_leaf | 52 |
| min_sum_hessian_in_leaf | 36 |
| max_bin | 404 |
| n_estimators | 14604 |
| learning_rate | 0.084236848 |
| max_depth | 31 |
| extra_trees | FALSE |
| path_smooth | 24 |
